# Antisense Oligonucleotide Quantification via Splint-Ligation PCR Assay in Non-Human Primate Central Nervous System Tissues and Biofluids

**DOI:** 10.1101/2025.01.19.630425

**Authors:** Thomas Jepp, Sarah Christian, Scott V. Dindot

## Abstract

Antisense oligonucleotides (ASOs) are chemically modified single-stranded oligonucleotides used to modulate the expression or processing of a target RNA transcript. The development of ASOs to treat human disease requires extensive preclinical studies in animal models. A critical component of these studies is determining the concentration of the ASO in tissues and biofluids, which are used to estimate the distribution, half-life, and dose-response relationship. The methods used to quantify ASOs are often constrained by low sensitivities, poor dynamic ranges, and the use of highly specialized equipment. Here, we describe the development of a Splint-Ligation-based quantitative PCR assay to measure the concentration of ASOs in nonhuman primate tissues and biofluids. Our results show that the Splint-Ligation Assay is highly sensitive and has a broad dynamic range in nonhuman primate central nervous system tissues and biofluids, ranging from picomolar to micromolar concentrations. Overall, our results show that the Splint-Ligation PCR Assay is a reliable, sensitive, and feasible method of ASO quantification.

## Introduction

Antisense oligonucleotides (ASOs) are chemically modified single-stranded oligonucleotides that modulate the expression of a gene upon hybridizing to a complementary RNA sequence. Phosphorothioate (PS) modification, the substitution of a non-bridging oxygen atom with a sulfur atom in phosphodiester bonds, is a near-ubiquitous feature of ASOs, increasing both the resistance of the molecule to degradation by nucleases [1, 2] and binding to plasma proteins [3, 4]. Modifications to the 2’-carbon of the ribose sugar, such as the locked nucleic acid (LNA) and 2’-O-Methoxyethyl (2’-MOE) modifications, increase both the resistance of the molecule to nucleases and binding affinity to the target sequence [5, 6]. Gapmer ASOs contain a ≥7nt long, unmodified DNA sequence (gap) flanked by chemically modified nucleosides on the 3’ and 5’ ends (wings). Gapmer ASOs repress the expression of a gene by inducing RNase H-mediated degradation of the RNA transcript [7]. Steric hindrance ASOs contain chemical modifications to all their nucleosides and therefore do not recruit RNase H, instead hindering the binding of molecules to the RNA sequence, often being used to redirect splicing [8, 9].

There is notable interest in developing ASOs to treat genetic disorders of the central nervous system (CNS) [10]. Preclinical nonhuman primate (NHP) studies of ASOs are a critical stage of research and development for novel therapies to characterize the pharmacological properties of investigational molecules in an animal closely related to humans. Results from these studies are used to determine the pharmacodynamic (PD) and pharmacokinetic (PK) properties of the ASO, which inform safety, dosages, and administration methods in subsequent human trials. PK studies evaluate the absorption, distribution, metabolism, and elimination of a drug and, therefore, rely on accurate measurements of ASO concentration in tissues and biofluids. Unfortunately, current standard methods of ASO concentration quantification in preclinical NHP CNS studies are often limited by some combination of poor sensitivity, narrow dynamic range, use of specialized equipment, the need to re-develop the assay for each ASO, and requiring large amounts of tissue. Due to the lack of an assay with sufficient sensitivity, dynamic range, and accessibility for the quantification of ASO concentration in CNS tissues, elements of valuable PK studies of this type in NHPs can be limited, for example, by lacking PK and PD data for all brain regions due to small sizes of samples [11], fewer brain regions being assessed [12], or lacking ASO concentration data [13].

Shin et al. recently developed a SplintR-Ligation PCR assay to quantify 2’-MOE-modified gapmer and steric hindrance ASOs in mouse serum and tissues [14]. Here, we developed a modified Splint-Ligation PCR assay to quantify the concentration of an LNA gapmer in NHP CNS tissues, plasma, and cerebrospinal fluid (CSF).

## Results

### Splint-Ligation Assay design

The Splint-Ligation Assay involves two oligonucleotide probes (Splint Probes 1 [57 nucleotides] and 2 [56 nucleotides]), corresponding to the 5’ and 3’ ends of the enhanced green fluorescent protein (eGFP) cDNA template sequence, plus 9-10 nucleotides of complementary sequence to the 5’ or 3’ end of the ASO (Figure S1). The Splint Probes are added to a sample containing the ASO, where the hybridization reaction is performed (Figure 1A). The Splint Probes hybridize to the ASO (Figure 1B), where they are ligated together by the SplintR ligase (Figure 1C). The concentration of the ligated probes, which is proportional to the ASO concentration of the sample, is then quantified by quantitative PCR (qPCR) using a commercially available eGFP TaqMan assay (Figure 1D). The TaqMan assay is specific to the Splint Probes, allowing the use of the same assay across ASOs.

**Figure 1.**
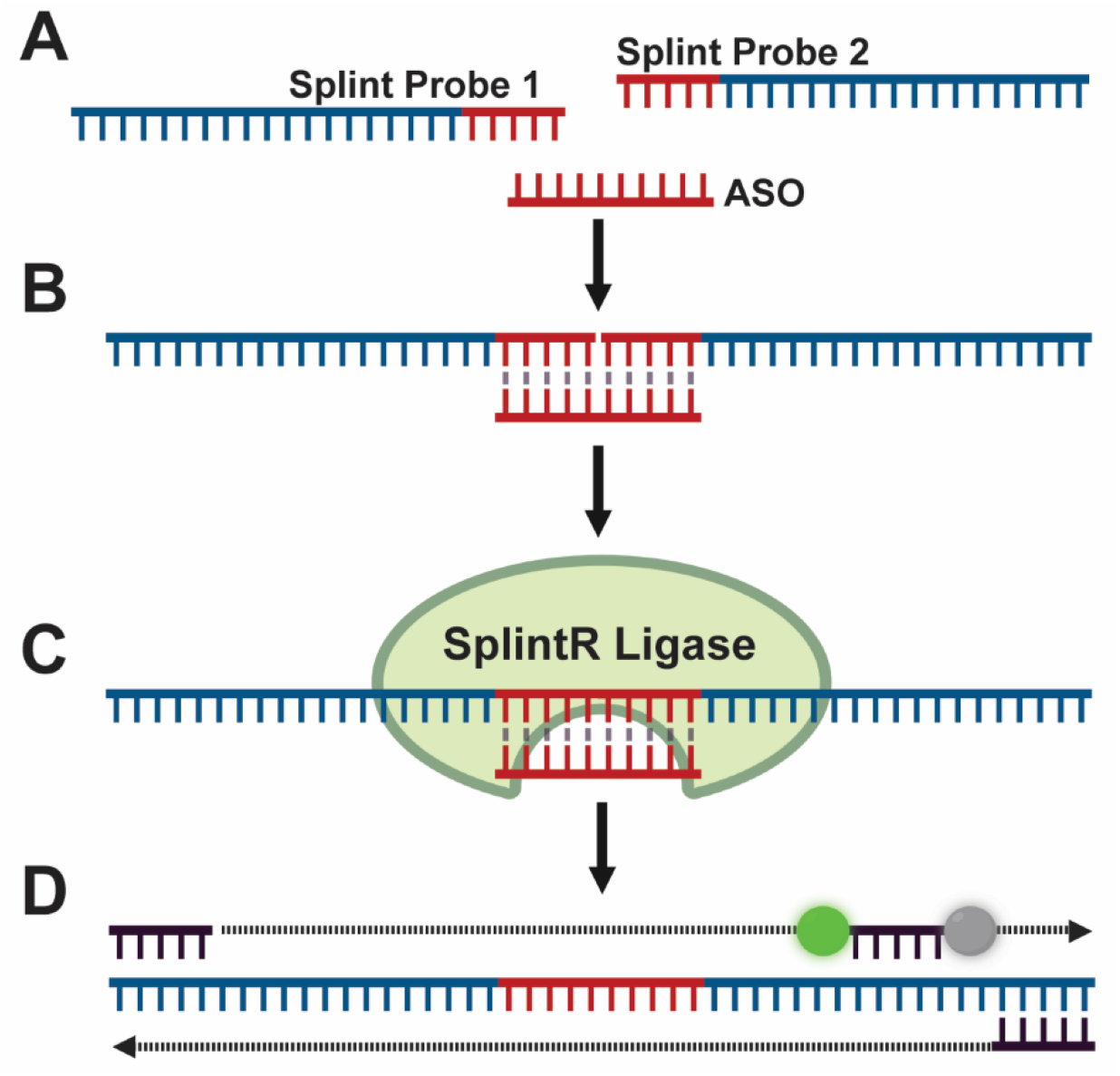
Schematic illustrating the Splint-Ligation PCR Assay for detection of ASO. **(A)** Both Splint Probes are added to processed sample. **(B)** A hybridization reaction is performed, hybridizing Splint Probes to the ASO. **(C)** SplintR Ligase is added to the mixture. A ligation reaction is performed, ligating splinted probes together. **(D)** An eGFP TaqMan assay is used to quantify the DNA ligation product.

### Splint-Ligation Assay in mouse brain tissue

To evaluate the utility of the Splint-Ligation Assay, we created an assay for an LNA gapmer ASO (18-mer, 3-11-4 wing-gap-wing, full PS backbone) from a previous study [15]. The assay was first tested using a dilution series of the LNA ASO in water (1 pM, 3.16 pM, 10 pM, 31.6 pM, 100 pM, 316 pM, 1 nM, 3.16 nM, 10 nM, 31.6 nM, 100 nM, 316 nM, 1 µM, 3.16 µM, and 10 µM; [*n* = 3 technical replicates]). There was a strong correlation (R^2^ = 0.992) between the ASO concentration and Cq values across the linear dynamic range (31.6 pM to 31.6 nM) (Figure 2A). A low level of ligated Splint Probes was detected (Cq = 29.22) in the negative control, indicating that the ligation reaction can occur without the ASO template.

**Figure 2.**
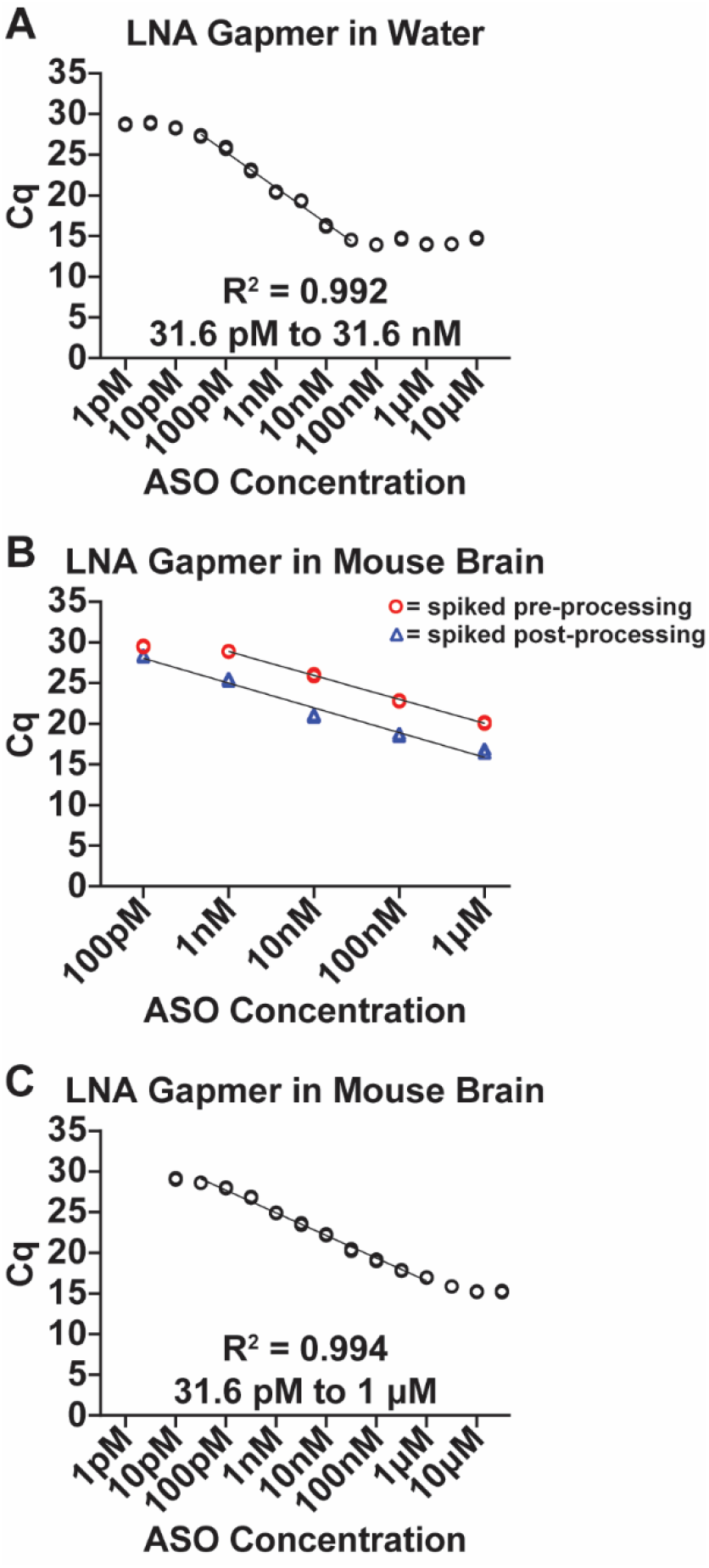
**(A)** Splint-Ligation quantification of a standard curve of the LNA gapmer ASO diluted in water (1 pM, 3.16 pM, 10 pM, 31.6 pM, 100 pM, 316 pM, 1 nM, 3.16 nM, 10 nM, 31.6 nM, 100 nM, 316 nM, 1 µM, 3.16 µM, 10µM; [*n* = 3 technical replicates]). **(B)** Splint-Ligation quantification of a standard curve of the LNA ASO in mouse brain homogenate (100 pM, 1 nM, 10 nM, 100 nM, 1 µM; [*n* = 3 technical replicates]), spiking ASO into sample before (R^2^ = 0.999, 1 nM to 1 µM) or after (R^2^ = 0.975, 100 pM to 1 µM) processing and removing cellular debris. Mean absolute difference between Cq values at each concentration: 100 pM = 1.1 Cq, 1 nM = 3.5 Cq, 10 nM = 5 Cq, 100 nM = 4.1 Cq, 1 µM = 3.5 Cq. 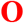 = spiked before, 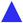 = spiked after. **(C)** Splint-Ligation quantification of a standard curve of the LNA ASO (10 pM, 31.6 pM, 100 pM, 316 pM, 1 nM, 3.16 nM, 10 nM, 31.6 nM, 100 nM, 316 nM, 1 µM, 3.16µM, 10 µM, 31.6 µM; [*n* =3 technical replicates]) in mouse whole brain lysate, spiked prior to homogenate incubation and centrifugation.

Standard curves are generated by spiking known concentrations of an ASO into control tissue homogenate. The control homogenate can be spiked with ASO before or after processing (i.e., incubation and centrifugation), and it is unclear whether some portion of the ASO is lost during processing. To test this, we performed the Splint-Ligation Assay on a dilution series of the LNA ASO (100 pM, 1 nM, 10 nM, 100 nM, and 1 µM; [*n* = 3 technical replicates]) in mouse brain homogenate under two conditions: adding ASO to the tissue 1) before or 2) after processing (see Methods). Cq values generated were significantly different between methods (mixed-effects linear regression, *P* < 0.001 [Figure 2B]), resulting in approximately 15-fold more ligated probe detected in the samples spiked after processing than those spiked before. These findings indicate that a portion of the ASO is lost during processing (e.g., by binding to cellular debris or insoluble proteins) and that spiking the control tissue homogenates after processing underestimates the ASO concentration of samples calculated using the standard curve.

To determine the dynamic range of the assay in the mouse brain, we performed the Splint-Ligation Assay on a broader dilution series of the LNA ASO (10 pM, 31.6 pM, 100 pM, 316 pM, 1 nM, 3.16 nM, 10 nM, 31.6 nM, 100 nM, 316 nM, 1 µM, 3.16 µM, 10 µM, and 31.6 µM; [*n* = 3 technical replicates]) in mouse whole brain homogenate, spiked before processing. There was a strong linear correlation (R^2^ = 0.994) between ASO concentration and the Cq values across a broad dynamic range (31.6 pM to 1 µM [Figure 2C]).

### Splint-Ligation Assay in NHP CNS tissues

Mammalian brains are comprised of regions that differ in cellular composition, cytoarchitecture, myelination, and extracellular matrix composition [16-19]. Preclinical studies of CNS-targeted ASOs in NHPs often assess the pharmacological properties of the molecule in different CNS regions [15]. We hypothesized that the efficiency of the Splint-Ligation Assay varies across CNS regions due to their different cellular and extracellular compositions. To test this, we compared the dynamic range, efficiency [slope of linear regression], and Cq value using a dilution series of the LNA ASO (10 pM, 100 pM, 1 nM, 10 nM, 100 nM, and 1 µM; [*n* = 3 technical replicates]) in 16 cynomolgus macaque CNS regions (cerebellum, cerebellar nuclei, caudate nucleus, cortical white matter, frontal cortex, globus pallidus, hippocampus, motor cortex, medulla, pons, putamen, thalamus, temporal lobe, cervical spinal cord, lumbar spinal cord, thoracic spinal cord [Figure S2]).

Comparing standard curves across 16 CNS regions requires PCR reactions on multiple 96-well plates. To examine the variability across plates (i.e., plate effect), the dynamic range, efficiency, Cq values, and percent coefficient of variation (%COV) of three standard curves each for motor and frontal cortex were compared. The splint-ligation reaction was repeated from the same serial dilutions each time, in order to account for variability from both this reaction and the qPCR. The efficiencies were similar and not significantly different (analysis of means with Nelson’s adjustment, α = 0.05 [Figure S3A]). The Cq values were not significantly different across the reactions and the plates for frontal cortex but were significantly different for motor cortex (mixed-effects linear regression: frontal cortex, *P* = 0.2; motor cortex, *P* <0.0001 [Figure S3A]). The percent coefficient of variation (%COV) of the Cq values was low across technical replicates per reaction (mean frontal cortex = 4.23%; mean motor cortex = 6.55%) but high across plates (mean frontal cortex = 15.1%; mean motor cortex = 54.21% [Table S1]). These results indicate the qPCR reaction can be variable across plates, and thus, reactions performed on different plates must be normalized to a standard curve from the same splint-ligation and qPCR reactions (see methods). The normalized Cq values were not significantly different across the reactions and plates (mixed-effects linear regression: frontal cortex, *P* = 0.9; motor cortex, *P* = 0.9 [Figure S3B]). The coefficient of variation (%COV) of the normalized Cq values was low across plates (mean frontal cortex = 8.84%; mean motor cortex = 9.53% [Table S2]).

Next, we compared the dynamic ranges, efficiencies, and normalized Cq values of the standard curves across the 16 CNS regions. The dynamic ranges were different among brain regions and reactions (Figure 3): 100 pM to 1 µM (cerebellar nuclei, cerebellum, cervical spinal cord, globus pallidus, lumbar spinal cord, medulla, thoracic spinal cord, temporal lobe) and 1 nM to 1 µM (caudate nuclei, cortical white matter, frontal cortex, hippocampus, motor cortex, pons, putamen, thalamus). The highest concentration (1 µM; 180,000 ng/g) examined was within the dynamic range, so the upper limit of quantification was not determined. Analysis of the overlapping dynamic range across regions (1nM to 1 µM) indicated that the efficiencies were not significantly different (analysis of means with Nelson’s adjustment, α = 0.05 [Figure S4]). In contrast, the normalized Cq values were significantly different (mixed-effects model, *P* < 0.0001) across the brain regions. Pairwise comparisons (Tukey’s HSD, α = 0.05) revealed 3 Tissue Groups that are similar and not significantly different (Table S3). The Tissue Groups consisted of the following CNS regions (Tissue group 1: frontal cortex, motor cortex, caudate nucleus, cortical white matter, hippocampus, medulla, pons, putamen, thalamus, temporal lobe; Tissue Group 2: globus pallidus, cervical spinal cord, thoracic spinal cord; Tissue Group 3: cerebellum, cerebellar nuclei [Table 1]). Collectively, these findings indicate that, although the CNS regions have similar efficiencies (slopes) and dynamic ranges, the Cq values of the curves are different between several regions. The CNS regions with similar efficiencies and Cq values can be assigned to groups and analyzed together via pooling control lysates and using this to make 1 standard curve. Regression analysis of Tissue Groups within their dynamic ranges revealed no significant difference in normalized Cq between tissues of the same Tissue Group (mixed-effect linear regression, [Tissue Group 1, *P* = 0.504; Tissue Group 2, *P* = 0.813; Tissue Group 3, *P* = 0.413]).

**Figure 3.**
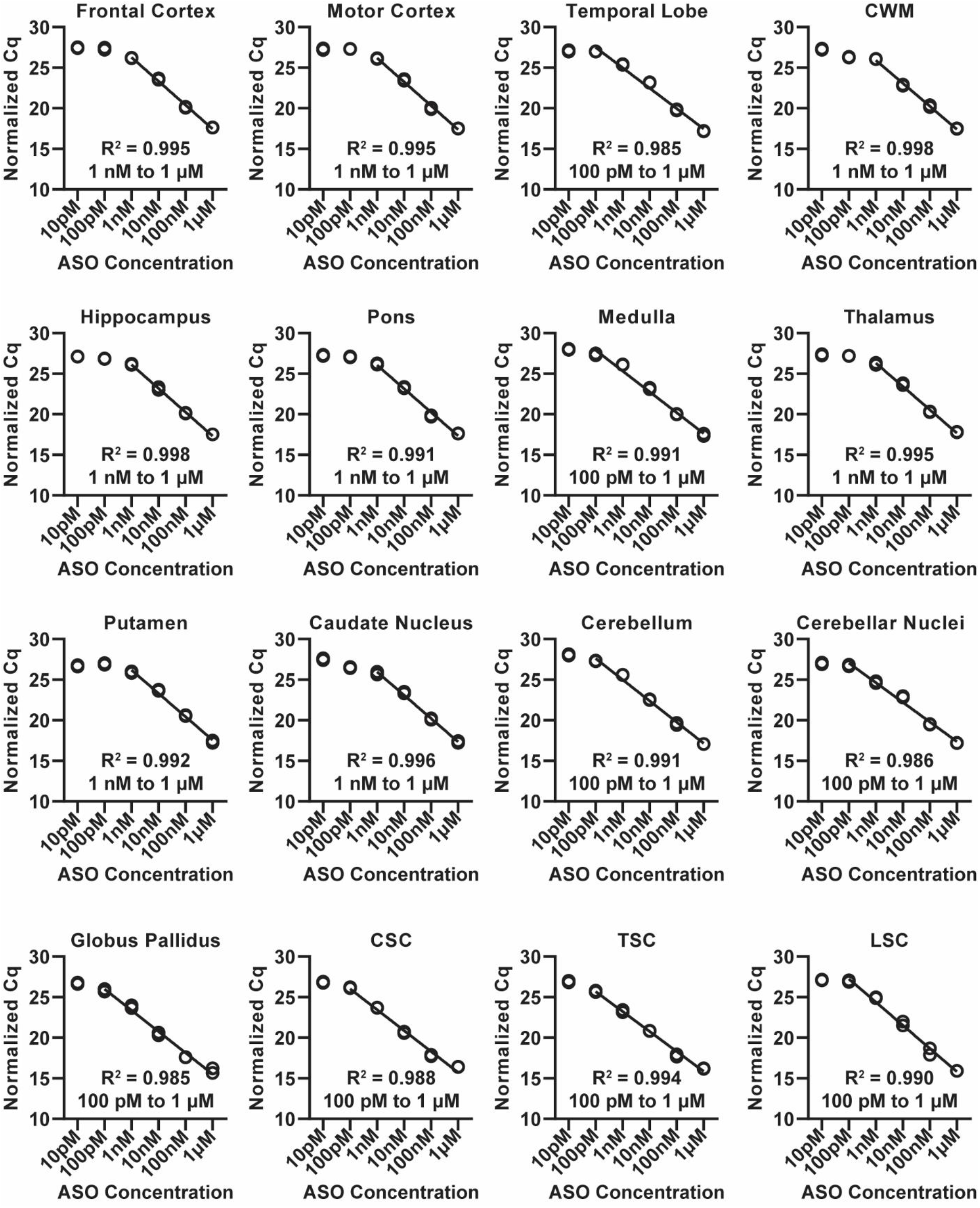
Splint-Ligation quantification of a dilution series of LNA ASO (10 pM, 100 pM, 1 nM, 10 nM, 100 nM, 1 µM; [*n* = 3 technical replicates]) in 16 CNS region tissues (cerebellum, cerebellar nuclei, caudate nucleus, cortical white matter, frontal cortex, globus pallidus, hippocampus, motor cortex, medulla, pons, putamen, thalamus, temporal lobe, cervical spinal cord, lumbar spinal cord, thoracic spinal cord) of a cynomolgus macaque. ASO concentration plotted against normalized Cq values. Abbreviations: CWM = cortical white matter, CSC = cervical spinal cord, TSC = thoracic spinal cord, LSC = lumbar spinal cord.

**Table 1.** NHP Tissue Group Dynamic Ranges.

| Table 1 – NHP Tissue Group Dynamic Ranges |  |  |  |
| --- | --- | --- | --- |
| Tissue | Tissue Group | Dynamic Range | Dynamic Range |
| Frontal Cortex | 1 | 1 nM to 1 μM | 180 to 180,000 ng/g |
| Motor Cortex |  |  |  |
| Caudate Nucleus |  |  |  |
| Cortical White Matter |  |  |  |
| Hippocampus |  |  |  |
| Medulla |  |  |  |
| Pons |  |  |  |
| Putamen |  |  |  |
| Thalamus |  |  |  |
| Temporal Lobe |  |  |  |
| Globus Pallidus | 2 | 100 pM to 1 μM | 18 to 180,000 ng/g |
| Cervical Spinal Cord |  |  |  |
| Thoracic Spinal Cord |  |  |  |
| Cerebellum | 3 | 100 pM to 1 μM | 18 to 180,000 ng/g |
| Cerebellar Nuclei |  |  |  |
| Lumbar Spinal Cord | N/A | 100 pM to 1 μM | 18 to 180,000 ng/g |

### Splint-Ligation Assay in NHP biofluids

To evaluate the Splint-Ligation Assay in NHP biofluids, standard curves of the LNA gapmer ASO (1 pM, 10 pM, 100 pM, 1 nM, 10 nM, 100 nM, and 1 µM; [*n* = 3 technical replicates]) were generated in cynomolgus plasma and CSF. Results showed that the assay had broad but different linear ranges in the samples (plasma: 1 pM to 1 µM, R^2^ = 0.994 [Figure 4A]; CSF: 100 pM to 1 µM, R^2^ = 0.984 [Figure 4B]).

**Figure 4.**
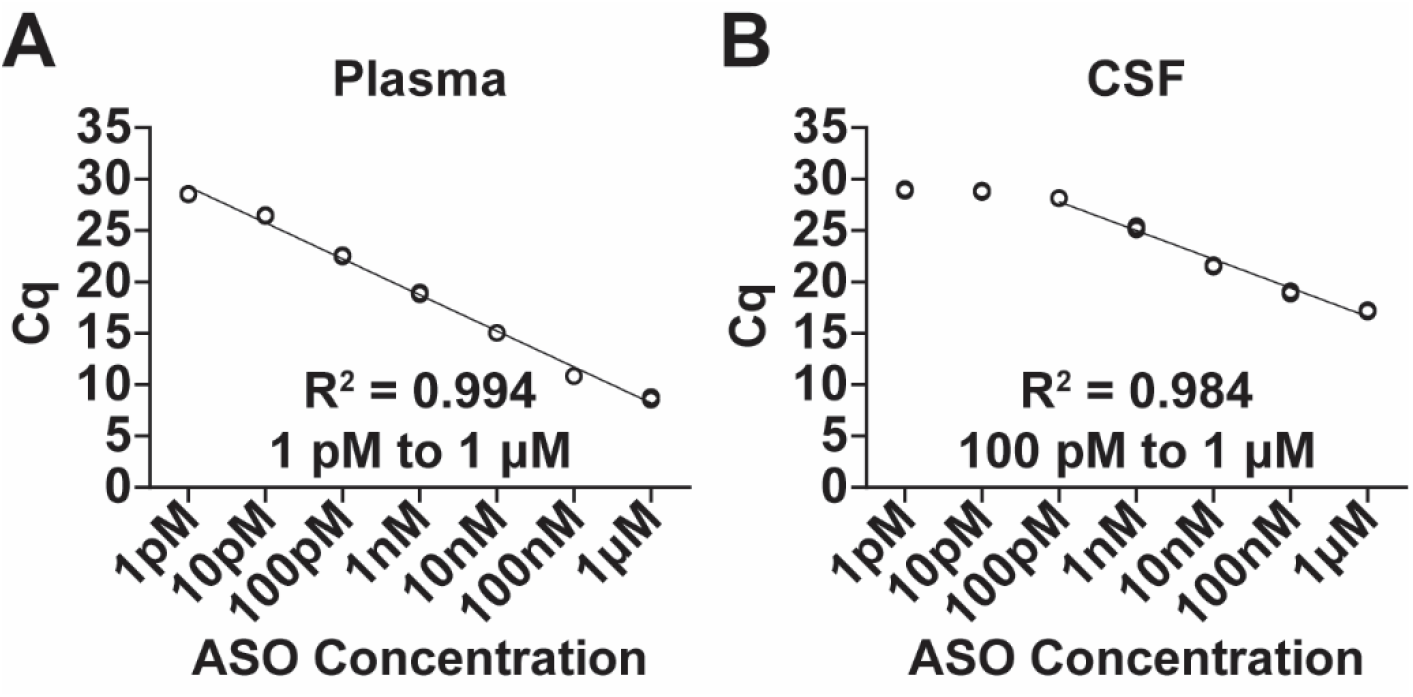
Splint-Ligation quantification of a dilution series of LNA ASO (1 pM, 10 pM, 100 pM, 1 nM, 10 nM, 100 nM, 1 µM; [*n* = 3 technical replicates]) in NHP **(A)** plasma and **(B)** CSF.

## Discussion

In this study, we show that the Splint-Ligation PCR Assay is a sensitive, simple, and accurate method for ASO quantification in NHP CNS tissues and biofluids. The design of the Splint Probes allows for using a single TaqMan assay to detect any ASO, reducing development time and complexity. We determined that controlling for matrix composition alone would not allow for accurate ASO quantification, thus the standard curves must be made in tissue homogenate before processing to avoid underestimation of sample ASO concentrations. We show that standard curves in 16 different NHP CNS tissues do not have a uniform relationship between ASO concentration and Cq value. However, we identified three groups of tissues with similar standard curves, allowing for the creation of standard curves in pooled control samples, minimizing the number of standard curves required to analyze different brain regions. Results from standard curves in NHP CNS tissues show a strong correlation between Cq values and ASO concentration within a linear dynamic range of up to 100 pM to 1 µM (18 to 180,000 ng/g). The Splint Ligation PCR Assay also demonstrated high sensitivity and a broad dynamic range in plasma (1 pM to 1 µM; 0.18 to 180,000 ng/mL) and CSF (100 pM to 1 µM; 18 to 180,000 ng/mL).

To control for the difference in matrix composition between samples and standards in ASO Splint-Ligation reactions, it was unclear whether ASO retrieval needed to be accounted for. Phosphorothioate ASOs bind to proteins [4], which make up a large proportion of the cellular debris that is removed from the samples by centrifugation during processing. We showed that a significant amount of the ASO is not retrieved when a sample is processed. By making serial dilutions via spiking the ASO into unprocessed homogenate immediately after tissue disruption and processing them in the same manner as the samples, we generate a standard curve that more accurately reflects the Cq values generated by a particular concentration of ASO in a sample.

For ethical and economic reasons, it is common to use relatively few control animals in NHP studies, often leading to a small number of control samples being available. We sought to mitigate the effect of this limit in the available control tissue matrix with which to make standard curves, as well as the time and resource burden imposed by running separate standard curves for every CNS region. By determining groups of tissues that can be run alongside each other with the same standard curves, we minimize tissue burden per control animal, reagents, labor, and plate space associated with running separate curves for every brain region, while ensuring that invalid comparisons aren’t made. We identified 3 Tissue Groups, within which there are no significant differences between tissues in Cq response to initial ASO concentration. As well as these broad groups, we show that it is possible to share standard curves between many smaller groups of tissues.

The assay is highly sensitive in NHP CNS samples, detecting down to 100 pM concentrations, approximately 18 ng of ASO per 1 g of tissue. This allows for less tissue to be used per sample, as balance sensitivity is more likely to constrain the mass of tissue required (<10 mg) than the sensitivity of the assay. It is, therefore, possible to use the same 4mm biopsy punch (∼40 mg) for both PD and PK assays, allowing accurate determination of the dose-response relationship by measuring protein, nucleic acid, and ASO concentrations from the same sample. Refining of reaction conditions and equipment used could likely lower the LLOQ (lower limit of quantification), and we expect that different ASO sequences and chemistries will have different dynamic ranges.

Drawbacks of previous methods of ASO quantification have been addressed by this assay. High-performance liquid chromatography and capillary gel electrophoresis methods used in PK studies have been shown to have LLOQs as high as 200 ng/mL in plasma and 600 ng/mL in tissues [20], 154 ng/mL, with a 25% COV in plasma [21], and 10 nM in plasma (standard curve 1: 10 nM to 500 nM, R^2^ = 0.962; standard curve 2: 0.5 µM to 20 µM, R^2^ = 0.998) [22]. Hybridization enzyme-linked immunosorbent assays (ELISAs) have achieved much greater sensitivities, with examples of LLOQs in tissues from studies including 1.52 ng/mL [23] and 0.77 ng/mL, with a 25% COV [21] in plasma. However, Hybridization ELISAs have limited dynamic ranges and require extensive sample processing. The Splint-Ligation PCR Assay has a low LLOQ and a broad dynamic range in plasma, with reliable detection demonstrated from 0.18 ng/mL to 180 µg/mL. The Splint-Ligation PCR Assay also has a low LLOQ and broad dynamic range in CNS tissue, reliably detecting as low as 18 to 180 ng/g and as high as 180 µg/g, with an average COV between 3 splint-ligation and qPCR reactions each of frontal and motor cortex of <10%.

In conclusion, we have developed a Splint-Ligation PCR Assay for the accurate quantification of ASOs in NHP CNS tissues and biofluids. The assay uses standard laboratory equipment and relatively inexpensive reagents, allowing it to be run in any lab with access to a qPCR thermocycler. The ability to design the Splint Probes around any available TaqMan assay, and only requiring a single TaqMan assay to detect the ASO, expedites the analysis of ASOs. The assay does not have to be redeveloped between ASOs, besides evaluating dynamic range. This method requires very small quantities of tissue (<10 mg) and does not compromise on the sensitivity of results compared to alternative methods. The assay also has a broad dynamic range and is therefore less likely to require dilutions and reruns, making it a very efficient method for assessing ASO concentration in limited samples. The use of this assay to determine PK parameters in preclinical studies could lead to a greater throughput of data, due to the increased speed, availability, and performance relative to other available methods, at a lower financial and time cost, leading to more efficient pharmacological studies for the treatment of genetic disorders of the CNS using ASOs.

## Materials and Methods

### Oligonucleotide synthesis

ASOs and Splint Probes were purchased from Integrated DNA Technologies (Coralville, IA) with a standard desalting purification method, and resuspended in nuclease free water (Invitrogen, Waltham, MA. Cat: 10977015). TaqMan Assay was purchased from Thermo Fisher Scientific (Waltham, MA), using sequences obtained from The Jackson Laboratory’s Protocol #22182 (Table S4).

### NHP use and sample collection

NHP CNS tissues were sampled from a cynomolgus macaque used as a control animal in a study performed by Charles River Laboratories (CRL; Montreal, Canada). The study was approved by the CR MTL Institutional Animal Care and Use Committee. During the study, the care and use of animals were conducted with guidance from the guidelines of the U.S. National Research Council and the Canadian Council on Animal Care. Following euthanasia, the brain was removed and cut coronally into 4-mm thick slices using a brain matrix. Tissue punches were taken from slices using 4mm biopsy punches corresponding to specific brain regions (Figure S2). The spinal cord was also removed, along with the dorsal root ganglia, and cut into cervical (C6 to C7), thoracic (T1 to T4), and lumbar (L6 and L7) sections. All samples were flash-frozen in liquid nitrogen before storing at -80°C. NHP biofluids were sampled from a cynomolgus macaque used as a control animal in a study performed by Shin Nippon Biomedical Laboratories (SNBL; Kagoshima, Japan). The study was approved by the SNBL Institutional Animal Care and Use Committee. During the study, the care and use of animals were conducted in accordance with the animal welfare bylaws of SNBL (AAALAC International accredited). CSF was collected from a lumbar catheter, placed into sterile tubes, and kept on ice. Samples were then centrifuged at 13,400 RCF at 2°C to 8°C for 1 minute, placed on dry ice until frozen, and stored at -80°C. Blood was drawn into a chilled sodium fluoride/potassium oxalate blood collection tube. These were then mixed by inversion 6 to 8 times and centrifuged at 1,000 RCF at 4°C for 10 minutes. Plasma was transferred into a new tube containing 5% cold phosphoric acid (1:10 [acid:plasma]), mixed by inversion at least 3 times, and frozen at -80°C, where they were stored.

### Sample and standard curve preparation

Assay Buffer was made up in nuclease free water (Invitrogen, Waltham, MA. Cat: 10977015), containing 10 mM Tris-HCl (pH 7.4), 150 mM NaCl, 1% NP-40, 0.5% sodium deoxycholate, 0.1% SDS, 5 mM EDTA, and 1 mM EGTA, and adjusted to pH 7.4. Tissue punches were cut into 2 to 4 roughly equal pieces, with 1 piece being retained for the use of this assay. Sample was transferred to a 2 mL locking lid microcentrifuge tube on dry ice, along with a 5mm stainless steel bead (Qiagen, Hilden, Germany. Cat: 69989). The mass of the sample was measured to within 0.1 mg. 3 µL of assay buffer was added to the tube, per 0.1 mg of tissue. Tubes were then disrupted via Tissuelyser II (Qiagen, Hilden, Germany. Cat: 85300) at 30hz for a total of 4 minutes (2x 2 minutes, rotating cassettes 180° between runs), repeating if full homogenization did not occur. Biofluids were not processed further, so standard curves were prepared in freshly thawed and well mixed CSF and plasma with no further steps.

Dilution series were generated by spiking the ASO into sample homogenate immediately after disruption at a concentration of 10 µM (or 10-fold higher than the greatest concentration in the standard curve). The spiked sample was then serially diluted 1:10 into more tissue homogenate until the lowest concentration required was reached. The spiked tissues were incubated on ice for 60 minutes, vortexing occasionally, then centrifuged at 16,000 RCF at 4°C for 30 minutes. The supernatant was collected and dispensed into a new microcentrifuge tube, then aliquoted and stored at -80°C.

### Ligation and hybridization reactions

Hybridization master mix was prepared. Per sample, this contains: 2 µL Splint-Probe 1 (100 nM), 2 µL Splint-Probe 2 (100 nM), 2 µL 10X SplintR Ligase buffer (New England Biolabs, Ipswich, MA. Cat: M0375), and 4 µL H_2_O. After mixing thoroughly, 10 µL of the hybridization master mix was pipetted into 0.2 mL PCR tubes. 2 µL of supernatant from each sample or standard was then also pipetted into the PCR tubes, mixed thoroughly by pipetting up and down 25 times at half the reaction volume (as vortexing leads to foaming), then briefly spun down.

This was placed in a thermocycler and heated to 95°C for 5 minutes, then cooled to 37°C at a rate of 0.1°C per second. The ligation master mix was then prepared. Per sample, 1 µL of SplintR Ligase Enzyme (New England Biolabs, Ipswich, MA. Cat: M0375) and 7 µL of H2O were added to a microcentrifuge tube, resulting in 25U of enzyme per reaction. This was then mixed, and 8 µL was added to each PCR tube, after the hybridization reaction was complete. The reaction was mixed thoroughly again by pipetting up and down as before, briefly spun down, heated to 37°C for 30 minutes, then heat inactivated at 65°C for 20 minutes.

### Quantitative PCR

Standard curves run on the same plate in the PCR reaction underwent the hybridization and ligation reactions in the same thermocycler run, using the same master mixes. PCR master mix was made up of 0.167 µL 60X TaqMan assay and 10 µL 2X TaqMan Gene Expression Master Mix (Applied Biosystems, Waltham MA. Cat: 4369016). 76.67 µL nuclease free water was added to each of the 20 µL products of the ligation reaction, mixed thoroughly by pipetting as before, then briefly spun down. 10.3 µL of PCR master mix was added to the wells of a 96-well, skirted PCR plate with solid white wells (Bio-Rad Laboratories, Hercules, CA. Cat: HSP9655), followed by 9.7 µL of the diluted standards in technical triplicates. The PCR plate was sealed with optically clear adhesive film (Bio-Rad Laboratories, Hercules, CA. Cat: MSB1001), briefly centrifuged, very gently mixed via vortexing, then briefly centrifuged again. qPCR reactions were run on a CFX96 Optical Reaction Module for Real-Time PCR Systems (Bio-Rad Laboratories, Hercules, CA. Cat: 1845097) mounted on a C1000 Touch Thermal Cycler Chassis (Bio-Rad Laboratories, Hercules, CA. Cat: 1841100), under the following conditions: 50°C for 2 minutes, 95°C for 10 minutes, 39x[95°C for 15s, 60°C for 1 minute, plate read]. Results were analyzed using CFX Maestro software (Bio-Rad Laboratories, Hercules, CA. Cat: 12013758).

### Statistical analysis

When investigating assay consistency across NHP CNS tissues, reactions were performed on four tissues at a time to ensure all reactions would fit on a single PCR plate. Each qPCR reaction’s relative fluorescence unit (RFU) threshold was set to a value that generated the highest average R^2^ for each of the four standard curves on the PCR plate. All Splint-Ligation PCR reactions contained either frontal cortex or motor cortex standard curves, made from the same dilution series. The average fold change in the Cq values of either motor or frontal cortex qPCR reactions, relative to the first reaction run, was calculated, and the normalization factor was applied to the Cq values of other tissues, to generate normalized Cq values. When determining average Cq for the frontal cortex and motor cortex, in order to determine the normalization factor for generating normalized Cq, outliers were determined as being the furthest replicate from the median, when technical replicates had a %COV of 2^Cq^ > 15%.

Statistical analyses were performed using JMP Pro 16 and GraphPad Prism 9. The dynamic ranges of all tissues were determined as the broadest concentration range with an R^2^ > 0.980 (mixed-effects linear regression, α = 0.05) between ASO concentration and Cq value. Statistical significance when comparing standard curves was performed by linear mixed-effect regression model; fixed effects used were tissue (or plate, when assessing plate effect), concentration, and interaction between the covariates (interaction was never found to be significant). Sample ID was included as a random variable, to account for repeated measures when performing technical replicates. Technical replicate variation was higher than desired, due to the nature of pipetting CNS homogenates and edge effects on PCR plates, necessitating outlier rejection on occasion. Reactions containing outlier replicates were determined by having a %COV of 2^Cq^ > 10%. In these cases, the Cq value furthest from the median was rejected. Rarely, technical replicate outliers were manually excluded (*n* = 4) or included (*n* = 2).

## Non-author contributors

We are grateful to Ashley Coffell, Tim Chiu, Luke Myers, Taira Saracco, and Alasdair Taylor for their valuable advice and critical feedback regarding this work. Figure 1 was created using BioRender: https://BioRender.com/v73t385.

## Author Contributions

**Tom Jepp**: conceptualization (equal); data curation (lead); formal analysis (equal); investigation (lead); methodology (equal); validation (lead); visualization (lead); writing – original draft (lead); writing – review & editing (equal). **Sarah Christian:** conceptualization (equal); funding acquisition (supporting); methodology (equal); project administration (equal); resources (equal); supervision (equal); writing – original draft (supporting); writing – review & editing (equal). **Scott V. Dindot:** conceptualization (equal); formal analysis (supporting); funding acquisition (lead); methodology (equal); project administration (lead); resources (equal); supervision (equal); visualization (supporting); writing – original draft (supporting); writing – review & editing (equal).

## References

[1] Matsukura M., Shinozuka K., Zon G., Mitsuya H., Reitz M., Cohen J. S., and Broder S. (1987). Phosphorothioate analogs of oligodeoxynucleotides: inhibitors of replication and cytopathic effects of human immunodeficiency virus. Proc Natl Acad Sci USA 84(21):7706–10

[2] Stein C. A., Subasinghe C., Shinozuka K., and Cohen J. S. (1988). Physicochemical properties of phosphorothioate oligodeoxynucleotides. Nucleic Acids Res 16(8):3209–21

[3] Crooke S. T., Vickers T. A., and Liang X. H. (2020). Phosphorothioate modified oligonucleotide-protein interactions. Nucleic Acids Res 48(10):5235–5253

[4] Liang X., Sun H., Shen W., and Crooke S. (2015). Identification and characterization of intracellular proteins that bind oligonucleotides with phosphorothioate linkages. Nucleic Acids Res 43(5):2927–2945

[5] Koshkin A., Nielsen P., Meldgaard M., Rajwanshi V., Singh S., and Wengel J. (1998). LNA (Locked Nucleic Acid): An RNA Mimic Forming Exceedingly Stable LNA:LNA Duplexes. J. Am. Chem. Soc. 120(50):13252–13253

[6] Altmann K. H., Fabbro D., Dean N. M., Geiger T., Monia B. P., Muller M., and Nicklin P. (1996). Second-generation antisense oligonucleotides: structure-activity relationships and the design of improved signal-transduction inhibitors. Biochem Soc Trans 24(3):630–7

[7] Monia B. P., Lesnik E. A., Gonzalez C., Lima W. F., McGee D., Guinosso C. J., Kawasaki A. M., Cook P. D., and Freier S. M. (1993). Evaluation of 2’-modified oligonucleotides containing 2’-deoxy gaps as antisense inhibitors of gene expression. J Biol Chem 268(19):14514–22

[8] Hua Y., Vickers T. A., Baker B. F., Bennett C. F., and Krainer A. R. (2007). Enhancement of SMN2 exon 7 inclusion by antisense oligonucleotides targeting the exon. PLoS Biol 5(4):e73

[9] Boiziau C., Kurfurst R., Cazenave C., Roig V., Thuong N. T., and Toulme J. J. (1991). Inhibition of translation initiation by antisense oligonucleotides via an RNase-H independent mechanism. Nucleic Acids Res 19(5):1113–9

[10] Bennett C., Kordasiewicz H., and Cleveland D. (2021). Antisense Drugs Make Sense for Neurological Diseases. Annu. Rev. Pharmacol. Toxicol. 61:831–852

[11] Jafar-Nejad P., Powers B., Soriano A., Zhao H., Norris D. A., Matson J., DeBrosse-Serra B., Watson J., Narayanan P., Chun S. J., Mazur C., … Rigo F. (2021). The atlas of RNase H antisense oligonucleotide distribution and activity in the CNS of rodents and non-human primates following central administration. Nucleic Acids Res 49(2):657–673

[12] Smith R. A., Miller T. M., Yamanaka K., Monia B. P., Condon T. P., Hung G., Lobsiger C. S., Ward C. M., McAlonis-Downes M., Wei H., Wancewicz E. V., … Cleveland D. W. (2006). Antisense oligonucleotide therapy for neurodegenerative disease. J Clin Invest 116(8):2290–6

[13] Kordasiewicz H. B., Stanek L. M., Wancewicz E. V., Mazur C., McAlonis M. M., Pytel K. A., Artates J. W., Weiss A., Cheng S. H., Shihabuddin L. S., Hung G., … Cleveland D. W. (2012). Sustained therapeutic reversal of Huntington’s disease by transient repression of huntingtin synthesis. Neuron 74(6):1031–44

[14] Shin M., Meda Krishnamurthy P., Devi G., and Watts J. K. (2022). Quantification of Antisense Oligonucleotides by Splint Ligation and Quantitative Polymerase Chain Reaction. Nucleic Acid Ther 32(1):66–73

[15] Dindot S. V., Christian S., Murphy W. J., Berent A., Panagoulias J., Schlafer A., Ballard J., Radeva K., Robinson R., Myers L., Jepp T., … Consortium F. (2023). An ASO therapy for Angelman syndrome that targets an evolutionarily conserved region at the start of the UBE3A-AS transcript. Sci Transl Med 15(688):eabf4077

[16] Dauth S., Grevesse T., Pantazopoulos H., Campbell P. H., Maoz B. M., Berretta S., and Parker K. K. (2016). Extracellular matrix protein expression is brain region dependent. J Comp Neurol 524(7):1309–36

[17] Brodmann K. Vergleichende Lokalisationslehre der Grophirnrinde [Comparative Localization Theory of the Cerebral Cortex], (1909) Johann Ambrosius Barth Verlag. Liepzig

[18] Mai J. K. and Majtanik M. (2023). Myeloarchitectonic maps of the human cerebral cortex registered to surface and sections of a standard atlas brain. Transl Neurosci 14(1):20220325

[19] Dauth S., Maoz B. M., Sheehy S. P., Hemphill M. A., Murty T., Macedonia M. K., Greer A. M., Budnik B., and Parker K. K. (2017). Neurons derived from different brain regions are inherently different in vitro: a novel multiregional brain-on-a-chip. J Neurophysiol 117(3):1320–1341

[20] Chen S. H., Qian M., Brennan J. M., and Gallo J. M. (1997). Determination of antisense phosphorothioate oligonucleotides and catabolites in biological fluids and tissue extracts using anion-exchange high-performance liquid chromatography and capillary gel electrophoresis. J Chromatogr B Biomed Sci Appl 692(1):43–51

[21] Sewell K. L., Geary R. S., Baker B. F., Glover J. M., Mant T. G., Yu R. Z., Tami J. A., and Dorr F. A. (2002). Phase I trial of ISIS 104838, a 2’-methoxyethyl modified antisense oligonucleotide targeting tumor necrosis factor-alpha. J Pharmacol Exp Ther 303(3):1334–43

[22] Leeds J. M., Graham M. J., Truong L., and Cummins L. L. (1996). Quantitation of phosphorothioate oligonucleotides in human plasma. Anal Biochem 235(1):36–43

[23] Yu R. Z., Kim T. W., Hong A., Watanabe T. A., Gaus H. J., and Geary R. S. (2007). Cross-species pharmacokinetic comparison from mouse to man of a second-generation antisense oligonucleotide, ISIS 301012, targeting human apolipoprotein B-100. Drug Metab Dispos 35(3):460–8

