## Supplementary Material for "Antisense Oligonucleotide Quantification via Splint-Ligation PCR Assay in Non-Human Primate Central Nervous System Tissues and Biofluids"

### Supplementary Figures

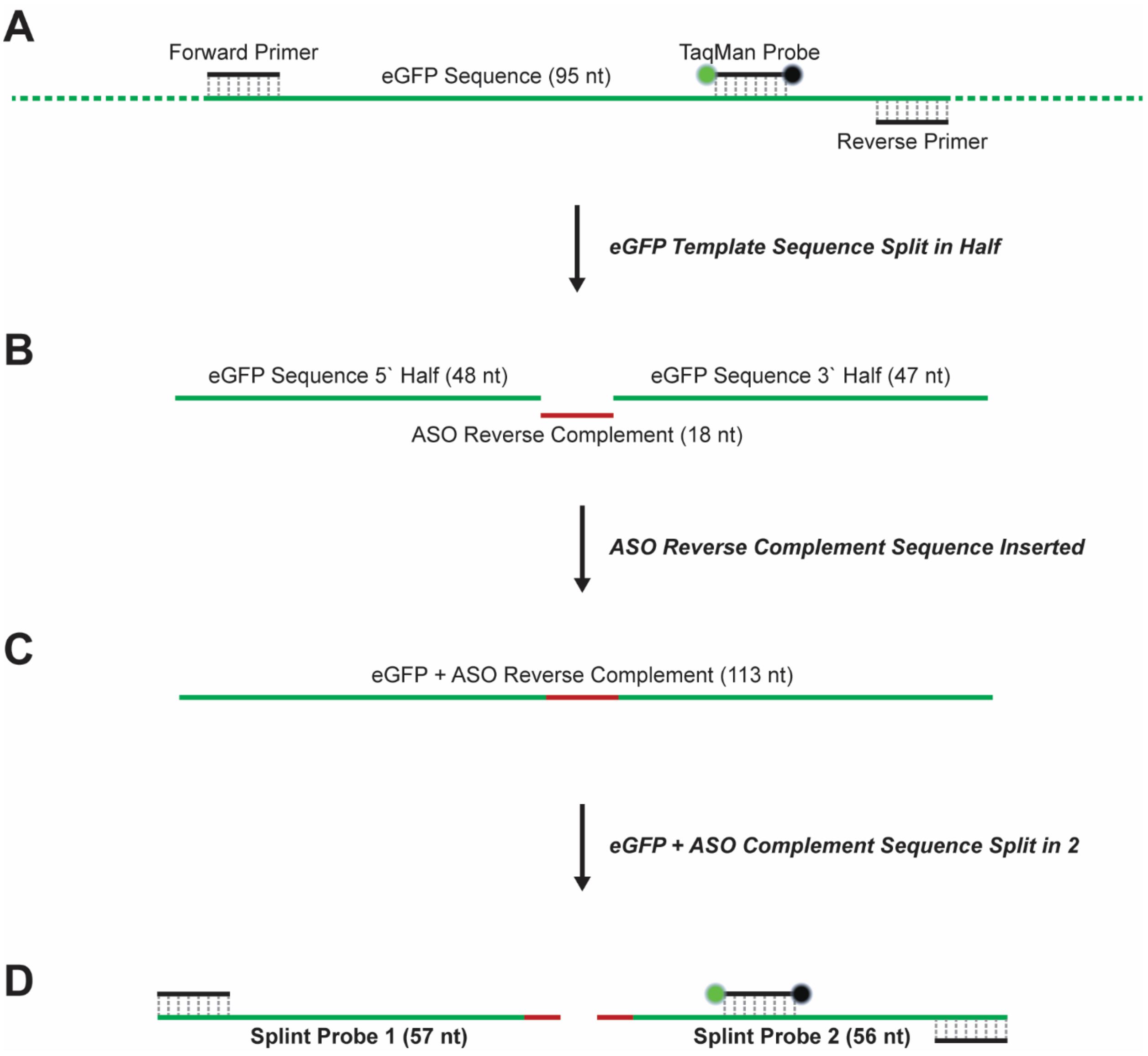


**Figure S1.** Schematic showing design of Splint Probes. **(A)** The template sequence of the TaqMan assay of interest is identified based on the location of the primers. **(B)** The template sequence is split into 2 roughly equal halves. **(C)** The ASO reverse complement sequence is added between the halves of the template sequence. **(D)** This template + ASO antisense sequence is split into 2 roughly equal halves, generating Splint Probe 1 and Splint Probe 2, which will be amplified by the TaqMan assay when ligated together.

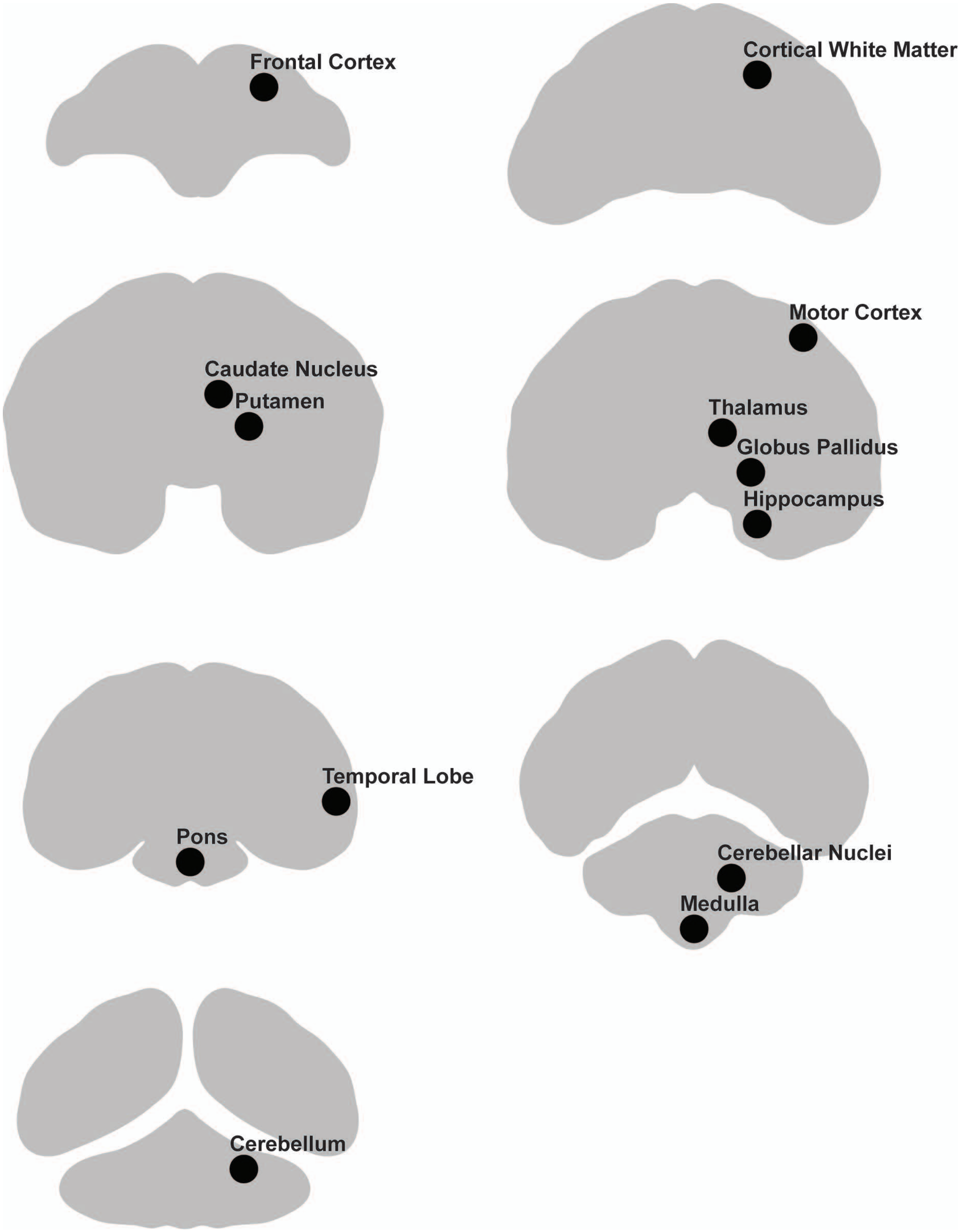


**Figure S2.** Schematic showing punch locations and slice number for NHP brain tissues. Slice order (left to right, top to bottom): 2, 4, 6, 8, 10, 12, 14.

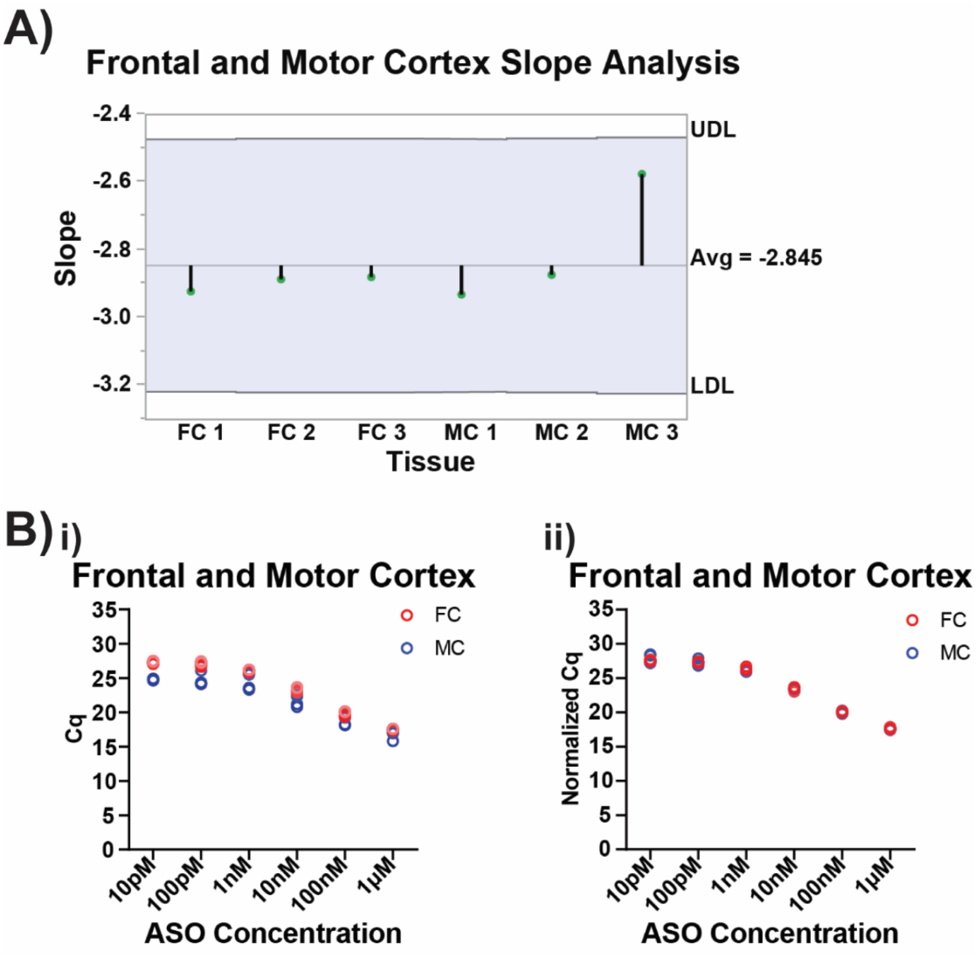


**Figure S3.** Splint-Ligation quantification of a dilution series of LNA ASO (10 pM, 100 pM, 1 nM, 10 nM, 100 nM, 1 µM; [*n* = 3 technical replicates]) in frontal cortex and motor cortex of a cynomolgus macaque. Splint-Ligation Assay was repeated 3 times in each tissue. Abbreviations: FC = frontal cortex, MC = motor cortex. **(A)** Slopes do not differ significantly between tissues or reactions (analysis of means with Nelson's adjustment, $\alpha = 0.05$). UDL = upper decision limit, LDL = lower decision limit, Avg = mean slope. **(B)** Results of replicates from each reaction. ASO concentration plotted against **(i)** Cq and **(ii)** Normalized Cq.

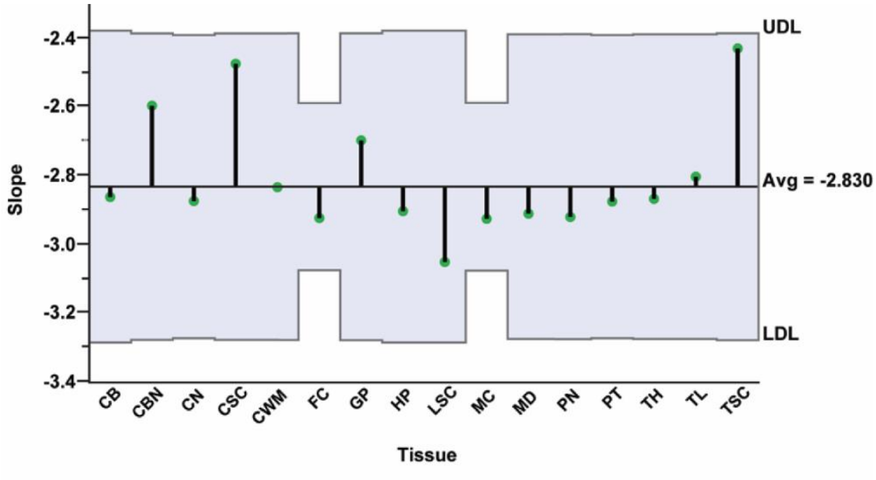


**Figure S4.** Analysis of means of Splint-Ligation PCR quantification of the slopes of standard curves of the LNA ASO (10 pM, 100 pM, 1 nM, 10 nM, 100 nM, 1 µM; [n = 3 technical replicates]) in 16 CNS region tissues (cerebellum, cerebellar nuclei, caudate nucleus, cortical white matter, frontal cortex, globus pallidus, hippocampus, motor cortex, medulla, pons, putamen, thalamus, temporal lobe, cervical spinal cord, lumbar spinal cord, thoracic spinal cord) of a cynomolgus macaque. Slopes do not differ significantly between tissues or reactions (analysis of means with Nelson's adjustment, $\alpha = 0.05$). Abbreviations: UDL = upper decision limit, LDL = lower decision limit, Avg = mean slope, CB = Cerebellum, CBN = Cerebellar Nuclei, CN = Caudate Nucleus, CSC = Cervical Spinal Cord, CWM = Cortical White Matter, FC = Frontal Cortex, GP = Globus Pallidus, HP = Hippocampus, LSC = Lumbar Spinal Cord, MC = Motor Cortex, MD = Medulla, PN = Pons, PT = Putamen, TH = Thalamus, TL = Temporal Lobe, TSC = Thoracic Spinal Cord.

### Supplementary Tables

**Table S1 - Cq variation**

| Tissue | Concentration | Plate 1 (Cq) | | | %COV | Plate 2 (Cq) | | | %COV | Plate 3 (Cq) | | | %COV | Total %COV |
|---|---|---|---|---|---|---|---|---|---|---|---|---|---|---|
| **Frontal Cortex** | 1 µM | | 17.61 | 17.63 | **0.98** | 17.59 | 17.53 | | **2.94** | | 17.5 | 17.35 | **7.35** | **6.92** |
| | 100 nM | 20.08 | 20.12 | 20.18 | **3.50** | 20.05 | 20.01 | | **1.96** | 19.38 | 19.58 | 19.54 | **7.20** | **20.37** |
| | 10 nM | 23.53 | 23.58 | 23.69 | **5.73** | 22.88 | 23.13 | 23.13 | **9.70** | 22.92 | 23.01 | 23.01 | **3.56** | **22.29** |
| | 1 nM | 26.24 | 26.17 | 26.19 | **2.51** | 26.16 | 26.19 | | **1.47** | 25.82 | 25.9 | | **3.92** | **10.84** |
| **Motor Cortex** | 1 µM | 17.51 | 17.54 | 17.54 | **1.20** | 17.01 | 16.98 | | **1.47** | 15.8 | | | | **35.83** |
| | 100 nM | 19.91 | 19.99 | 20.08 | **5.90** | 19.22 | 19.39 | | **8.32** | 18.11 | 18.23 | | **5.88** | **47.55** |
| | 10 nM | 23.39 | 23.47 | 23.6 | **7.40** | 22.34 | 22.52 | | **8.81** | 20.8 | 21.2 | | **19.48** | **61.81** |
| | 1 nM | 26.17 | 26.1 | | **3.43** | 25.54 | 25.52 | | **0.98** | 23.42 | 23.58 | 23.32 | **9.17** | **71.64** |

**Table S2 - Normalized Cq variation**

| Tissue | Concentration | Plate 1 (Normalized Cq) | | | %COV | Plate 2 (Normalized Cq) | | | %COV | Plate 3 (Normalized Cq) | | | %COV | Total %COV |
|---|---|---|---|---|---|---|---|---|---|---|---|---|---|---|
| **Frontal Cortex** | 1 µM | 17.61 | 17.63 | | **0.98** | 17.7 | 17.64 | | **2.94** | 17.85 | 17.7 | | **7.35** | **6.26** |
| | 100 nM | 20.08 | 20.12 | 20.18 | **3.50** | 20.17 | 20.13 | | **1.96** | 19.77 | 19.98 | 19.94 | **7.57** | **9.35** |
| | 10 nM | 23.53 | 23.58 | 23.69 | **5.73** | 23.02 | 23.27 | 23.27 | **9.70** | 23.38 | 23.47 | 23.47 | **3.56** | **13.40** |
| | 1 nM | 26.24 | 26.17 | 26.19 | **2.51** | 26.32 | 26.35 | | **1.47** | 26.34 | 26.42 | | 3.92 | **6.36** |
| **Motor Cortex** | 1 µM | 17.51 | 17.54 | 17.54 | **1.20** | 17.61 | 17.58 | | **1.47** | 17.57 | | | | **2.46** |
| | 100 nM | 19.91 | 19.99 | 20.08 | **5.90** | 19.9 | 20.08 | | **8.81** | 20.13 | 20.27 | | **6.86** | **9.19** |
| | 10 nM | 23.39 | 23.47 | 23.6 | **7.40** | 23.13 | 23.32 | | **9.30** | 23.12 | 23.57 | | **21.88** | **13.22** |
| | 1 nM | 26.17 | 26.1 | | **3.43** | 26.44 | 26.42 | | **0.98** | 26.04 | 26.22 | 25.93 | **10.25** | **13.24** |

| Table S3 – Pairwise Normalized Cq Comparisons (Tukey's HSD) | | |
|---|---|---|
| **Tissue 1** | **Tissue 2** | **Prob>\|t\|** |
| Cerebellum | Cerebellar Nuclei | 1.0000 |
| Cerebellum | Caudate Nucleus | 0.7983 |
| Cerebellum | Cervical Spinal Cord | <.0001 |
| Cerebellum | Cortical White Matter | 0.7870 |
| Cerebellum | Frontal Cortex | 0.0636 |
| Cerebellum | Globus Pallidus | <.0001 |
| Cerebellum | Hippocampus | 0.5705 |
| Cerebellum | Lumbar Spinal Cord | 0.0137 |
| Cerebellum | Motor Cortex | 0.1493 |
| Cerebellum | Medulla | 0.7237 |
| Cerebellum | Pons | 0.6197 |
| Cerebellum | Putamen | 0.2196 |
| Cerebellum | Thalamus | 0.0656 |
| Cerebellum | Temporal Lobe | 0.9999 |
| Cerebellum | Thoracic Spinal Cord | <.0001 |
| Cerebellar Nuclei | Caudate Nucleus | 0.4580 |
| Cerebellar Nuclei | Cervical Spinal Cord | <.0001 |
| Cerebellar Nuclei | Cortical White Matter | 0.4462 |
| Cerebellar Nuclei | Frontal Cortex | 0.0106 |
| Cerebellar Nuclei | Globus Pallidus | <.0001 |
| Cerebellar Nuclei | Hippocampus | 0.2548 |
| Cerebellar Nuclei | Lumbar Spinal Cord | 0.0501 |
| Cerebellar Nuclei | Motor Cortex | 0.0300 |
| Cerebellar Nuclei | Medulla | 0.3784 |
| Cerebellar Nuclei | Pons | 0.2892 |
| Cerebellar Nuclei | Putamen | 0.0672 |
| Cerebellar Nuclei | Thalamus | 0.0158 |
| Cerebellar Nuclei | Temporal Lobe | 0.9891 |
| Cerebellar Nuclei | Thoracic Spinal Cord | <.0001 |
| Caudate Nucleus | Cervical Spinal Cord | <.0001 |
| Caudate Nucleus | Cortical White Matter | 1.0000 |
| Caudate Nucleus | Frontal Cortex | 0.9993 |
| Caudate Nucleus | Globus Pallidus | <.0001 |
| Caudate Nucleus | Hippocampus | 1.0000 |
| Caudate Nucleus | Lumbar Spinal Cord | <.0001 |
| Caudate Nucleus | Motor Cortex | 1.0000 |
| Caudate Nucleus | Medulla | 1.0000 |
| Caudate Nucleus | Pons | 1.0000 |
| Caudate Nucleus | Putamen | 0.9998 |

| | | |
|---|---|---|
| Caudate Nucleus | Thalamus | 0.9778 |
| Caudate Nucleus | Temporal Lobe | 0.9982 |
| Caudate Nucleus | Thoracic Spinal Cord | <.0001 |
| Cervical Spinal Cord | Cortical White Matter | <.0001 |
| Cervical Spinal Cord | Frontal Cortex | <.0001 |
| Cervical Spinal Cord | Globus Pallidus | 1.0000 |
| Cervical Spinal Cord | Hippocampus | <.0001 |
| Cervical Spinal Cord | Lumbar Spinal Cord | 0.4756 |
| Cervical Spinal Cord | Motor Cortex | <.0001 |
| Cervical Spinal Cord | Medulla | <.0001 |
| Cervical Spinal Cord | Pons | <.0001 |
| Cervical Spinal Cord | Putamen | <.0001 |
| Cervical Spinal Cord | Thalamus | <.0001 |
| Cervical Spinal Cord | Temporal Lobe | <.0001 |
| Cervical Spinal Cord | Thoracic Spinal Cord | 1.0000 |
| Cortical White Matter | Frontal Cortex | 0.9996 |
| Cortical White Matter | Globus Pallidus | <.0001 |
| Cortical White Matter | Hippocampus | 1.0000 |
| Cortical White Matter | Lumbar Spinal Cord | <.0001 |
| Cortical White Matter | Motor Cortex | 1.0000 |
| Cortical White Matter | Medulla | 1.0000 |
| Cortical White Matter | Pons | 1.0000 |
| Cortical White Matter | Putamen | 0.9999 |
| Cortical White Matter | Thalamus | 0.9821 |
| Cortical White Matter | Temporal Lobe | 0.9977 |
| Cortical White Matter | Thoracic Spinal Cord | <.0001 |
| Frontal Cortex | Globus Pallidus | <.0001 |
| Frontal Cortex | Hippocampus | 1.0000 |
| Frontal Cortex | Lumbar Spinal Cord | <.0001 |
| Frontal Cortex | Motor Cortex | 1.0000 |
| Frontal Cortex | Medulla | 0.9999 |
| Frontal Cortex | Pons | 1.0000 |
| Frontal Cortex | Putamen | 1.0000 |
| Frontal Cortex | Thalamus | 1.0000 |
| Frontal Cortex | Temporal Lobe | 0.5001 |
| Frontal Cortex | Thoracic Spinal Cord | <.0001 |
| Globus Pallidus | Hippocampus | <.0001 |
| Globus Pallidus | Lumbar Spinal Cord | 0.1685 |
| Globus Pallidus | Motor Cortex | <.0001 |
| Globus Pallidus | Medulla | <.0001 |
| Globus Pallidus | Pons | <.0001 |

| | | |
|---|---|---|
| Globus Pallidus | Putamen | <.0001 |
| Globus Pallidus | Thalamus | <.0001 |
| Globus Pallidus | Temporal Lobe | <.0001 |
| Globus Pallidus | Thoracic Spinal Cord | 1.0000 |
| Hippocampus | Lumbar Spinal Cord | <.0001 |
| Hippocampus | Motor Cortex | 1.0000 |
| Hippocampus | Medulla | 1.0000 |
| Hippocampus | Pons | 1.0000 |
| Hippocampus | Putamen | 1.0000 |
| Hippocampus | Thalamus | 0.9989 |
| Hippocampus | Temporal Lobe | 0.9737 |
| Hippocampus | Thoracic Spinal Cord | <.0001 |
| Lumbar Spinal Cord | Motor Cortex | <.0001 |
| Lumbar Spinal Cord | Medulla | <.0001 |
| Lumbar Spinal Cord | Pons | <.0001 |
| Lumbar Spinal Cord | Putamen | <.0001 |
| Lumbar Spinal Cord | Thalamus | <.0001 |
| Lumbar Spinal Cord | Temporal Lobe | 0.0008 |
| Lumbar Spinal Cord | Thoracic Spinal Cord | 0.2263 |
| Motor Cortex | Medulla | 1.0000 |
| Motor Cortex | Pons | 1.0000 |
| Motor Cortex | Putamen | 1.0000 |
| Motor Cortex | Thalamus | 0.9987 |
| Motor Cortex | Temporal Lobe | 0.7453 |
| Motor Cortex | Thoracic Spinal Cord | <.0001 |
| Medulla | Pons | 1.0000 |
| Medulla | Putamen | 1.0000 |
| Medulla | Thalamus | 0.9905 |
| Medulla | Temporal Lobe | 0.9946 |
| Medulla | Thoracic Spinal Cord | <.0001 |
| Pons | Putamen | 1.0000 |
| Pons | Thalamus | 0.9974 |
| Pons | Temporal Lobe | 0.9834 |
| Pons | Thoracic Spinal Cord | <.0001 |
| Putamen | Thalamus | 1.0000 |
| Putamen | Temporal Lobe | 0.7480 |
| Putamen | Thoracic Spinal Cord | <.0001 |
| Thalamus | Temporal Lobe | 0.3889 |
| Thalamus | Thoracic Spinal Cord | <.0001 |
| Temporal Lobe | Thoracic Spinal Cord | <.0001 |

| Table S4 – Oligonucleotide Sequences | |
|---|---|
| **Oligonucleotide** | **Sequence (5’-3’)** |
| LNA gapmer | +A*+G*+A*A*T*G*G*C*A*C*A*T*C*T*+C*+T*+T*+G |
| 5’ Splint Probe | AAGGGCATCGACTTCAAGGAGGACGGCAACATCCTGGGGCACAAGCTGCAAGAGATG |
| 3’ Splint Probe | /5Phos/TGCCATTCTGAGTACAACTACAACAGCCACAACGTCTATATCATGGCCGACAAGCA |
| TaqMan Forward Primer | AAGGGCATCGACTTCAAGG |
| TaqMan Reverse Primer | TGCTTGTCGGCCATGATATAG |
| TaqMan Probe | FAM-CTTGTGCCCCAGGATGTTGCC-Quencher |
| | + = LNA, * = PS, /5Phos/ = 5’ Phosphorylation |
